# Identification of TIA1 mRNA targets during human neuronal development

**DOI:** 10.1101/2021.01.26.428265

**Authors:** Loryn P. Byres, Marat Mufteev, Kyoko E. Yuki, Wei Wei, Alina Piekna, Michael D. Wilson, Deivid C. Rodrigues, James Ellis

## Abstract

**Background:** Neuronal development is a tightly controlled process involving multi-layered regulatory mechanisms. While transcriptional pathways regulating neurodevelopment are well characterized, post-transcriptional programs are still poorly understood. TIA1 is an RNA-binding protein that can regulate splicing, stability, or translation of target mRNAs, and has been shown to play critical roles in neurodevelopment. However, the identity of mRNAs regulated by TIA1 during neurodevelopment is still unknown.

**Methods and Results:** To identify the mRNAs targeted by TIA1 during the first stages of human neurodevelopment, we performed RNA immunoprecipitation-sequencing (RIP-seq) on human embryonic stem cells (hESCs) and derived neural progenitor cells (NPCs), and cortical neurons. While there was no change in TIA1 protein levels, the number of TIA1 targeted mRNAs decreased from pluripotent cells to neurons. We identified 2400, 845, and 330 TIA1 mRNA targets in hESCs, NPC, and neurons, respectively. The vast majority of mRNA targets in hESC were genes associated with neurodevelopment and included autism spectrum disorder-risk genes that were not bound in neurons. Additionally, we found that most TIA1 mRNA targets have reduced ribosomal engagement levels.

**Conclusion:** Our results reveal TIA1 mRNA targets in hESCs and during human neurodevelopment, indicate that translation repression is a key process targeted by TIA1 binding and implicate TIA1 function in neuronal differentiation.

## Introduction

Post-transcriptional regulation is the collection of mechanisms regulating gene expression that start operating co-transcriptionally through the processing of the pre-mRNAs, and plays critical roles in cellular homeostasis and development [1]. Post-transcriptional mechanisms contribute to the differential gene expression profile of cells containing the same genetic blueprint. They modify the proteome allowing for the manifestation of distinct physiological properties in a cell-specific manner, independent of transcriptional regulation [2, 3]. RNA-binding proteins (RBPs) are modular regulatory proteins containing one or more domains for specific contact with RNAs [4, 5]. RBPs play a central role in post-transcriptional regulation through their association with mRNAs in all stages of mRNA metabolism, from mRNA biogenesis in the nucleus to sub-cellular localization and degradation in the cytoplasm [6].

Post-transcriptional regulation mechanisms play pivotal roles during neurodevelopment and in neuronal function [7–9]. Importantly, abnormal post-transcriptional regulation associates with neurological diseases and is consistently implicated in neurodevelopmental disorders including autism spectrum disorder (ASD). For example, mutations in multiple RBP genes have been shown to increase the risk for ASD [10].

TIA1 is an RBP that recruits RNAs into stress granules as part of the adaptive response to stress [11, 12]. However, while efforts have been made to understand TIA1 function in the stress response supporting cell survival, it is becoming clear that TIA1 also contributes to cellular physiology in homeostatic states. TIA1 interacts with the splicing machinery facilitating alternative splicing site recognition [13, 14]. It can also regulate translation of mRNAs during steady-state conditions or during development after binding to AU-rich or U-rich elements present in target mRNAs [15–17]. TIA1 functions can vary in a cell-type specific manner and independent of stress states. For example, knockdown of TIA1 protein levels can either promote or repress cell proliferation depending on the cell type [18].

Immunoprecipitation of TIA1 followed by transcriptome-wide analyses of co-eluted RNAs have been carried out on colorectal carcinoma (RKO) and HEK293 cells for unbiased identification of TIA1 target RNAs in non-stressed conditions [19, 20]. In these studies, TIA1 binds a variety of mRNAs exerting a range of cellular functions, including transcriptional regulation, metabolism, and cell cycle progression. TIA1 has also been shown to have important roles in neuronal development, synaptic plasticity, and memory in mice [17, 21]. However, a comprehensive list of target transcripts, and the underlying role of TIA1 in early development and neuronal physiology, have yet to be uncovered.

Here we performed RNA immunoprecipitation (RIP) of TIA1 protein followed by next-generation sequencing of co-eluted RNAs (RIP-seq) using an *in vitro* model of neurodevelopment involving differentiation from human embryonic stem cells (hESCs), to neural progenitor cells (NPCs) and then to cortical neurons. To precisely map and quantify mRNAs bound by TIA1 at the 3ʹUTR isoform level, we performed 3ʹUTR RNA-seq on transcripts precipitated by a TIA1 specific antibody and normalized these to RNAs eluted with an isogenic IgG control. We found thousands of high confidence target mRNAs in hESCs. Interestingly, the number of targets and their binding efficiencies decrease significantly in NPCs and neurons, indicating that the TIA1 regulatory network of target mRNAs decreases as hESCs differentiate into neurons. Most TIA1 mRNA targets have reduced ribosomal engagement levels, suggesting that translational repression throughout the first stages of neurodevelopment is a key process associated with TIA1 binding.

## Results

### RIP-seq to discover TIA1 mRNA targets during neurodevelopment

To identify TIA1 mRNA targets transcriptome-wide in an *in vitro* model of neurodevelopment, we conducted RIP-seq on samples collected from each stage of neuronal differentiation as reported previously [17] (Figure 1). Three replicates of pluripotent hESCs (H9 line), and their derived NPCs and cortical neurons were used for RNA extraction and immunoprecipitation of TIA1 (IP). Given that TIA1 binding sites are enriched at the 3ʹUTR of target mRNAs [20, 22], for quantification of co-eluted mRNAs we deployed 3ʹUTR mRNA profiling using an automated QuantSeq protocol (see methods). This approach precisely maps the polyadenylation sites and quantifies mRNA at the 3ʹUTR isoform level [23]. Each cell lysate was divided into 3 aliquots: input RNA sample, TIA1 IP sample, and IgG IP sample as the negative control (Figure 1A). The specificity of the TIA1 antibody used was determined by western blots that showed reduction in levels of a protein with expected TIA1 size (43 kDa) following shRNA-mediated TIA1 mRNA knock-down experiments (Figure 1B). Recovery of TIA1 protein after IP was also confirmed by western blot (Figure 1C). The presence of *MECP2* and *UBE3A* mRNAs, expressed in all cell types and known TIA1 mRNA targets with validated significance for neurodevelopment and neurodevelopmental disorders [17, 24], were evaluated by qRT-PCR in hESC IP samples. Figure 1D shows that, as expected by successful co-elution of specific mRNA targets, *MECP2* and *UBE3A* mRNAs were enriched in TIA1 IP but not in IgG IP samples. 18S RNA was used as negative RNA control.

**Figure 1.**
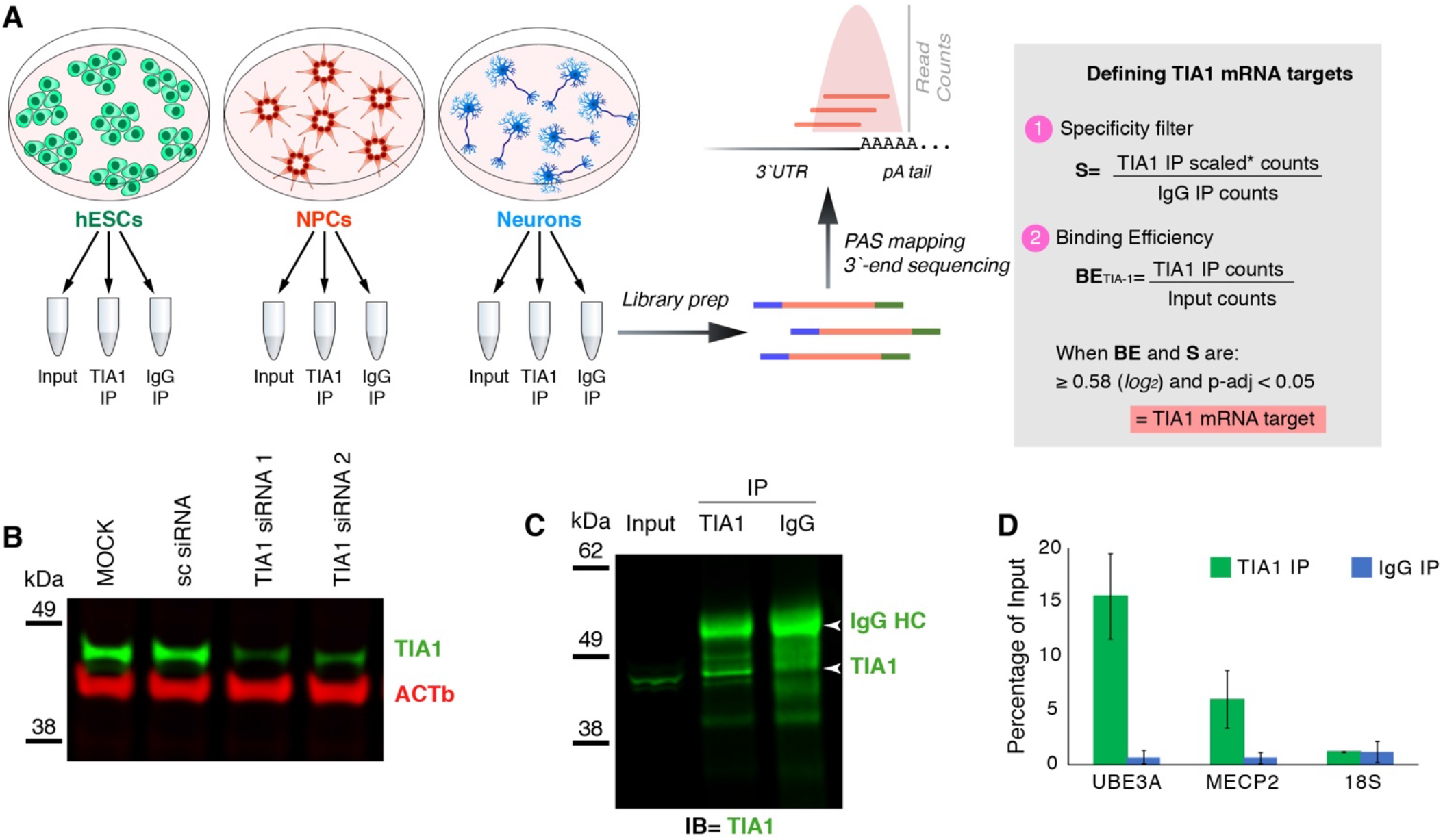
Experimental set up and validations. A, schematic representation of the experimental set up. hESCs, and derived NPCs and cortical neurons were used to collect total RNA (input), TIA1 and IgG-bound RNAs (TIA1-IP and IgG IP). Three replicates of each cell type were used for downstream experiments. Collected RNAs were processed for library preparation and sequenced using 3ʹ-end based sequencing technique, followed by PolyAdenylation Site (PAS) mapping and quantification of transcripts. TIA1 target mRNAs were defined by calculating the ratios of TIA1 over Input and IgG IP’s samples. When the TIA1 enrichment was 1.5-fold (or 0.58x in log_2_ scale) higher in both Input and IgG with adjusted p-value ≤ 0.05, the mRNA was called a TIA1 target. B-D, The TIA1 antibody used for IP was validated by western blots and RIP-qRT-PCR. B, western blot shows decrease of TIA1 protein levels upon shRNA-based knock-down of TIA1 mRNAs. C, western blot of IP samples show detection of TIA1 and IgG heavy chain (HC) proteins at expected sizes. D, RIP-qRT-PCR shows that MECP2 and UBE3A transcripts were co-eluted in hESCs TIA1 IP samples. 18S RNA was used as negative control. * please refer to methods for scaling procedure.

Overall quality of total RNA used as input samples from all aliquots and cell types was determined by bioanalyzer analysis, which showed that all input samples had RNA integrity numbers (RIN) values greater than 7 (Supplementary Figure 1A). Quality of RNA samples from TIA1 or IgG IPs could not be determined by bioanalyzer since these samples lack the ribosomal RNA molecules used to calculate RIN values. Nonetheless, analyses of RNA length showed that the fraction of RNAs larger than 200 nucleotides (nt) was above 90% in all TIA1 IP samples, indicating that the majority of RNA molecules remained intact after immunoprecipitation and RNA purification steps (Supplementary Figure 1B). In all three independent biological replicates for each cell type the total RNA recovered from IgG IP samples was significantly lower than the TIA1 IP samples. Two of the IgG samples, one replicate each of NPCs and neurons, did not have sufficient RNA material for library prep despite the high number of cells used and our efforts to collect all possible co-eluted RNA (Supplementary Figure 1C). This resulted in the use of two replicates of IgG IP samples for NPC and neuron cell types for downstream analyses. The low recovery of RNAs from IgG IP samples indicates that our IP method resulted in low background of non-specific RNAs bound to the magnetic beads.

After sequencing, we confirmed the cellular identities using cell-type gene-specific markers from the input samples (Figure 2A). High reproducibility of the datasets was confirmed by principle component analysis (PCA) on the entire RIP-seq dataset, which showed significant separation between cell types (Figure 2B). We also observed significant separation between IP and input samples in the same cell type indicating that TIA1 co-eluted a fraction of specific mRNAs different from the IgG fraction (Figure 2C).

**Figure 2.**
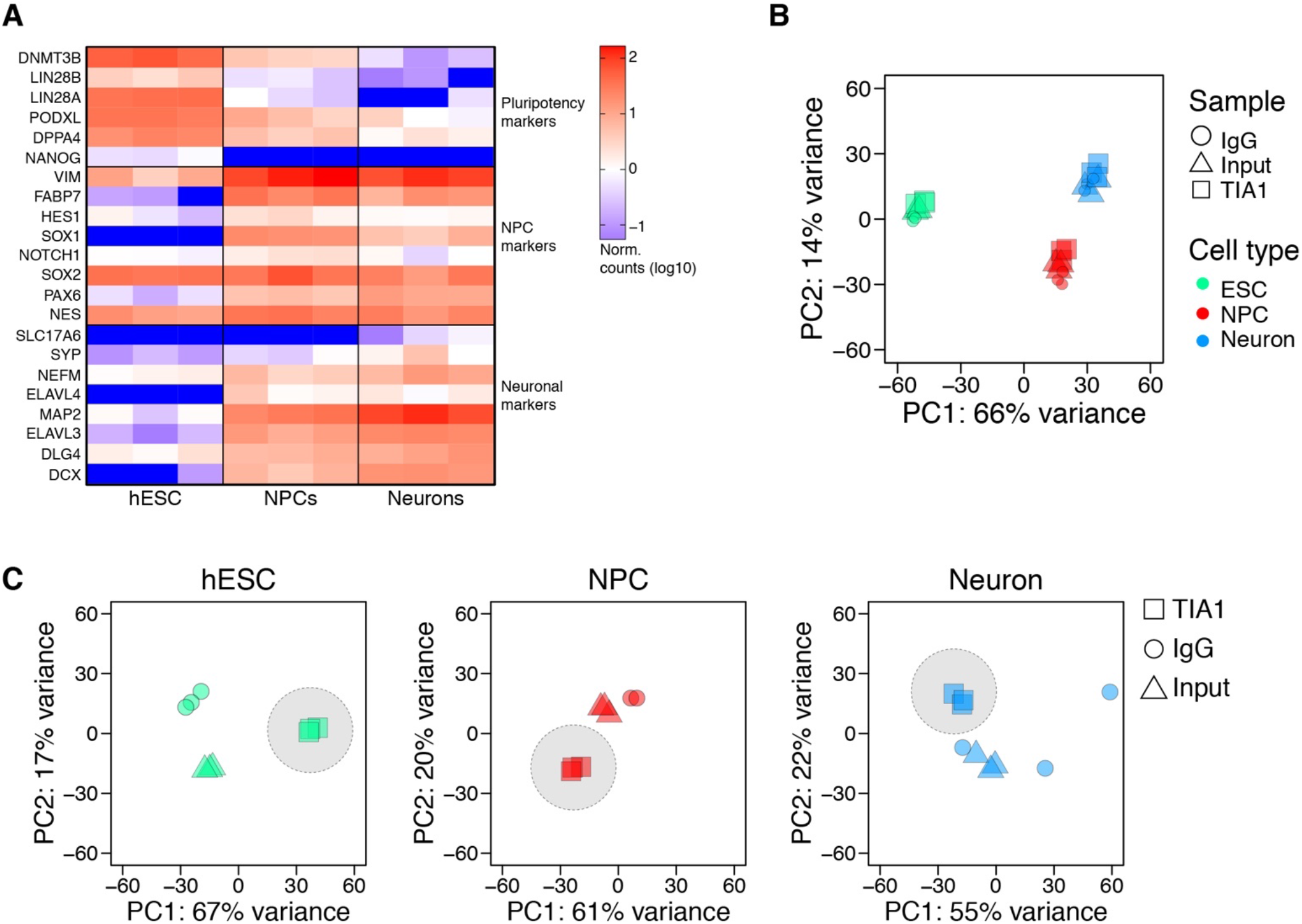
Confirmation of cell types and quality control of replicates. A, Analysis of read counts normalized to medians of each replicate and cell-type (log_10_ scale) showing abundance of cell-type specific gene markers. B, principal component analysis showing significant separation of samples according to cell type. C, principal component analysis showing separation of samples for each cell type according to the sample type.

### Identification of TIA1 targets during neurodevelopment

To identify TIA1 mRNA targets we first calculated TIA1 binding efficiency (BE) by determining the ratios of all detected mRNAs in IP over the input samples [12, 25] (Figure 1A, and methods). The average BE and its accuracy for each transcript across all three biological replicates was determined using DESeq2 [26]. To reduce noise artifacts caused by IP, we selected target mRNAs with high enrichment of TIA1 over IgG IP (Figure 1A). This method identified 2,400 mRNAs in hESCs, 845 mRNAs in NPCs, and 330 mRNAs in neurons as TIA1 targets, representing 18%, 7%, and 3% of the total expressed transcriptome in the respective cell types (Supplementary table 1). Reassuringly, the known TIA1 target mRNAs *TP53* [27], *MECP2* [17], and *UBE3A* [20] were successfully identified as TIA1 targets in hESCs with similar BE values (Figure 3A, and supplementary table 1). In contrast, mRNAs encoding ribosomal proteins, which have not been identified as TIA1 targets in existing CLIP-seq datasets from other cell types [12, 13, 20] had significantly lower TIA1 BEs relative to other mRNAs in hESCs (Figure 3B).

**Figure 3.**
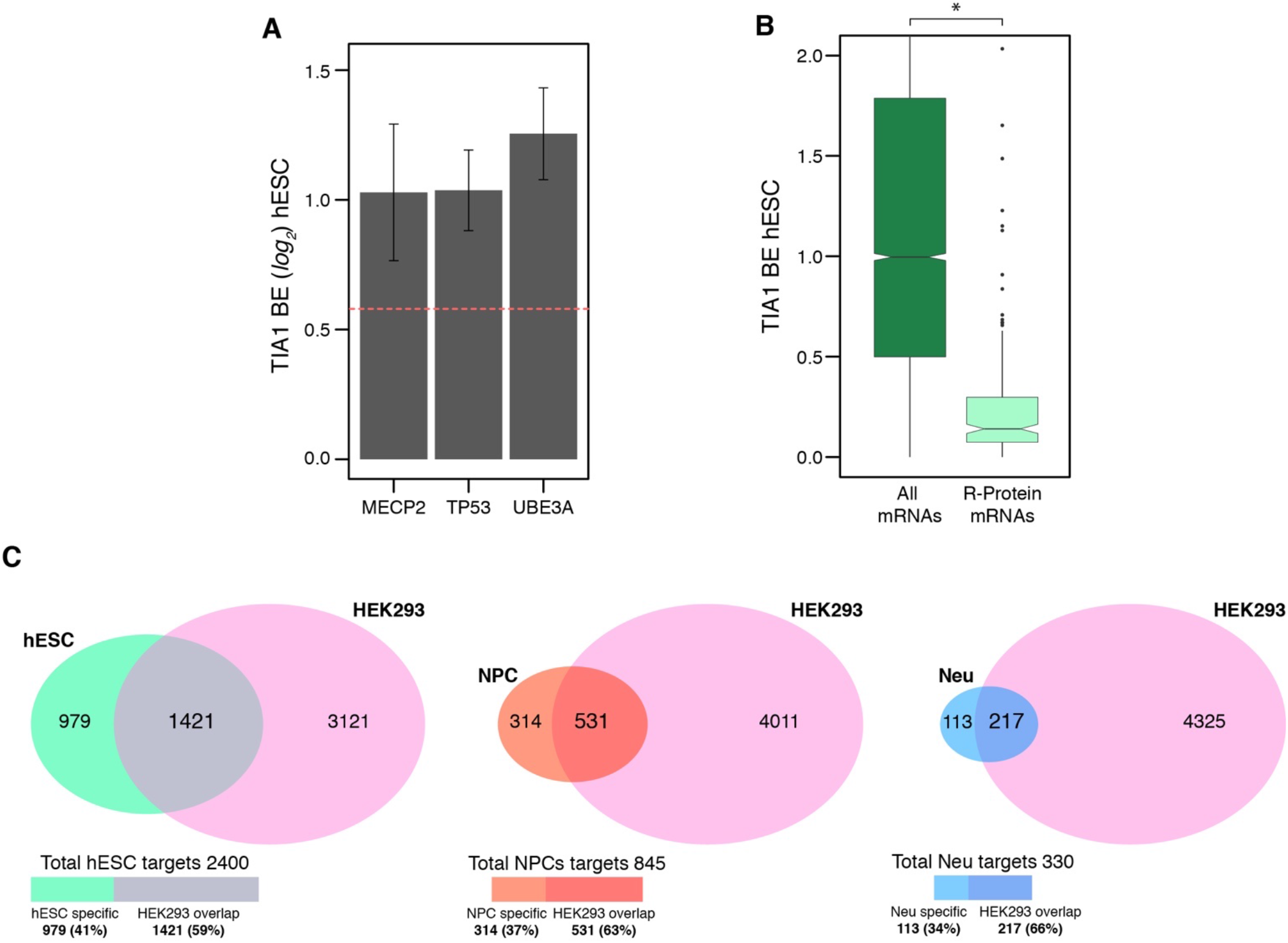
Defining TIA1 BE and discovery of target mRNAs. A, TIA1 binding efficiency (BE) of three previously known TIA1 mRNA targets: TP53, MECP2, and UBE3A in hESCs. The threshold for positive calling of TIA1 binding is above 0.58 (log_2_ scale – red dashed line). B, TIA1 binding efficiency of mRNAs encoding ribosomal proteins (R-Proteins), in comparison to all detected transcripts. *p < 0.05 (two-tailed student’s t-test). C, Comparison of our dataset with previously published CLIP-seq dataset of TIA1 target mRNAs for all cell types.

Previous CLIP-seq experiments had identified 5,411 TIA1 mRNA targets in HEK293 cells [20]. Approximately 84% of these targets were found to be expressed in at least one of the cell types in our neurodevelopmental model (Supplementary table 1). Of these, we found that between approximately 60 to 65% of TIA1 targets identified in our cell types are consistent with HEK293-ascertained targets (Figure 3C and Supplementary table 1).

To gain insight into the processes regulated by TIA1 binding, we performed gene enrichment analysis of TIA1 targeted mRNAs from cell types. TIA1 targets were enriched for terms associated with cell cycle progression, metabolism, and transcriptional regulation (Supplementary table 2). Interestingly, in hESCs we identified an enrichment for terms associated with development and cellular differentiation (Figure 4A). Examination of that cluster revealed that it was further enriched for terms associated with cellular differentiation of multiple tissues and cell types from different embryonic origins such as neuronal, mesenchyme, muscle, and hematopoietic tissues (Figure 4B), indicating that these key cell-fate determination genes are expressed at the pluripotent stage and regulated by TIA1. Further examination of the “nervous system development” gene ontology set showed that it included not only critical genes for neurodevelopment, but also genes associated with high-risk for ASD (e.g. *AFF2* and *CHD8*), genes implicated in neuronal syndromes (*USP9X*), and genes found to be translationally dysregulated in Rett syndrome cortical neurons (NEDD4L) (Figure 4C and Supplementary table 2).

**Figure 4.**
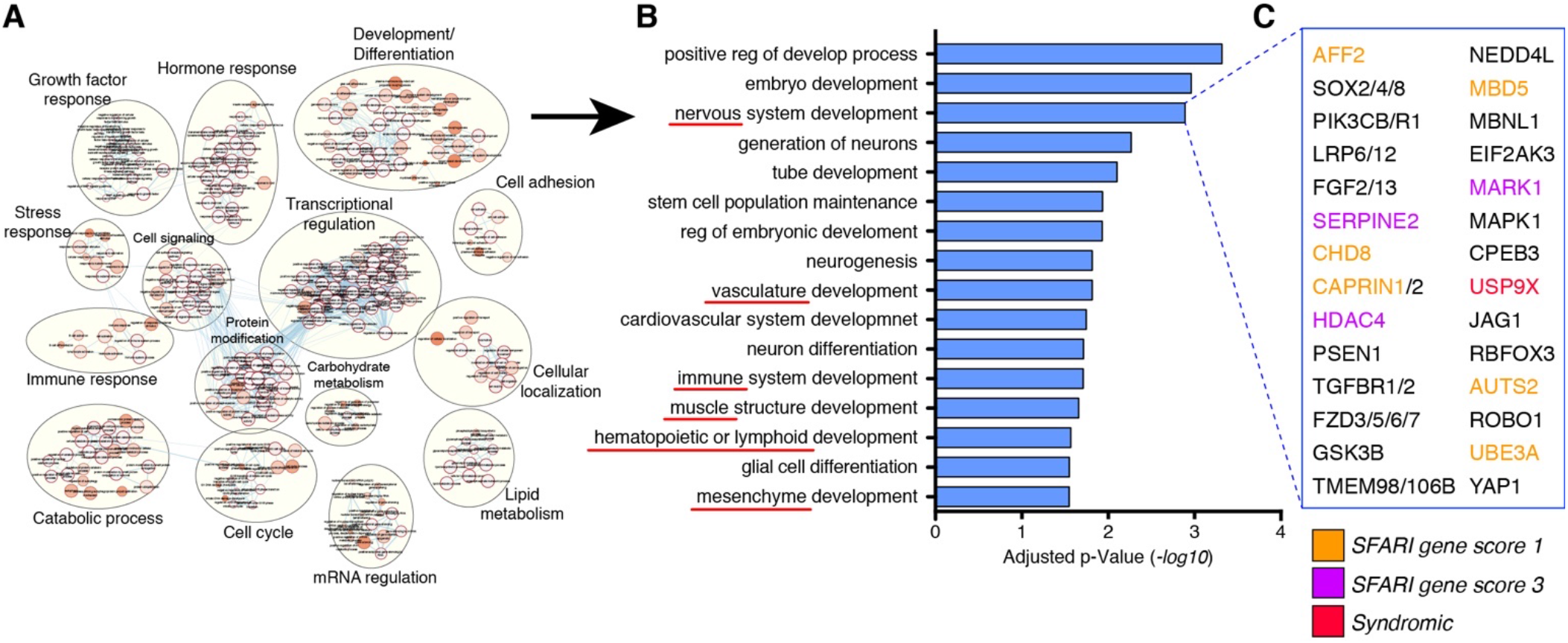
Functional analysis of RIP-seq dataset. A, Gene ontology analysis show enrichment of developmental processes associated genes as TIA1 target mRNAs in hESCs. B, example of gene ontologies enriched in “development and differentiation” processes. Red underlines indicate processes related to development into tissues from diverse embryonic origins that are expressed in hESCs and bound by TIA1. C, example of genes enriched in “nervous system development” with critical roles in neurodevelopment, ASD-associated (orange and pink), and syndromic-associate (red).

To compare TIA1 binding between cell types, and focus on transcripts where changes in TIA1 binding may contribute to neurodevelopment, we analysed target mRNAs in hESCs that were still expressed in the other cell types. We detected 2,290 TIA1 target mRNAs in hESCs that were also expressed in NPCs and neurons. However, the number of TIA1 target mRNAs decreased significantly to 836 in NPCs, and 325 in neurons (Figure 5A-B). Most of the NPC and neuronal targets were also bound by TIA1 in hESCs (Figure 5A). The decrease in number of TIA1 targets during neurodevelopment was confirmed to be due to a systematic decrease in BE of the hESC-specific TIA1 targeted mRNAs in NPCs and neurons relative to hESCs (Figure 5B-C). Interestingly, when including targets detected in all cells, we found that TIA1 BE values were also overall reduced in NPCs and neuron (Figure 5D). Together, these results indicate that TIA1 regulation through mRNA binding is reduced as neurodevelopment progresses.

We also detected 143 ASD-risk genes (SFARI gene score ≥3) as TIA1 mRNA targets that are consistently expressed across all cell types (129, 54, and 17 targets detected in hESCs, NPCs, and neurons, respectively - Supplementary Table 3). Interestingly, of these, 80.4% are bound by TIA1 in either hESCs, or in both hESCs and NPCs but not in neurons (Supplementary Table 3). Only 17 ASD-risk genes were bound by TIA1 in neurons, accounting for 11.9% of all TIA1 ASD-risk gene targets, and 14 of which are consistently bound by TIA1 in all three cell types. Together, our data suggests that TIA1 regulates neurodevelopment and ASD-relevant transcripts in the early stages of development, and this regulation is reduced as cells progress through neurodevelopment.

**Figure 5.**
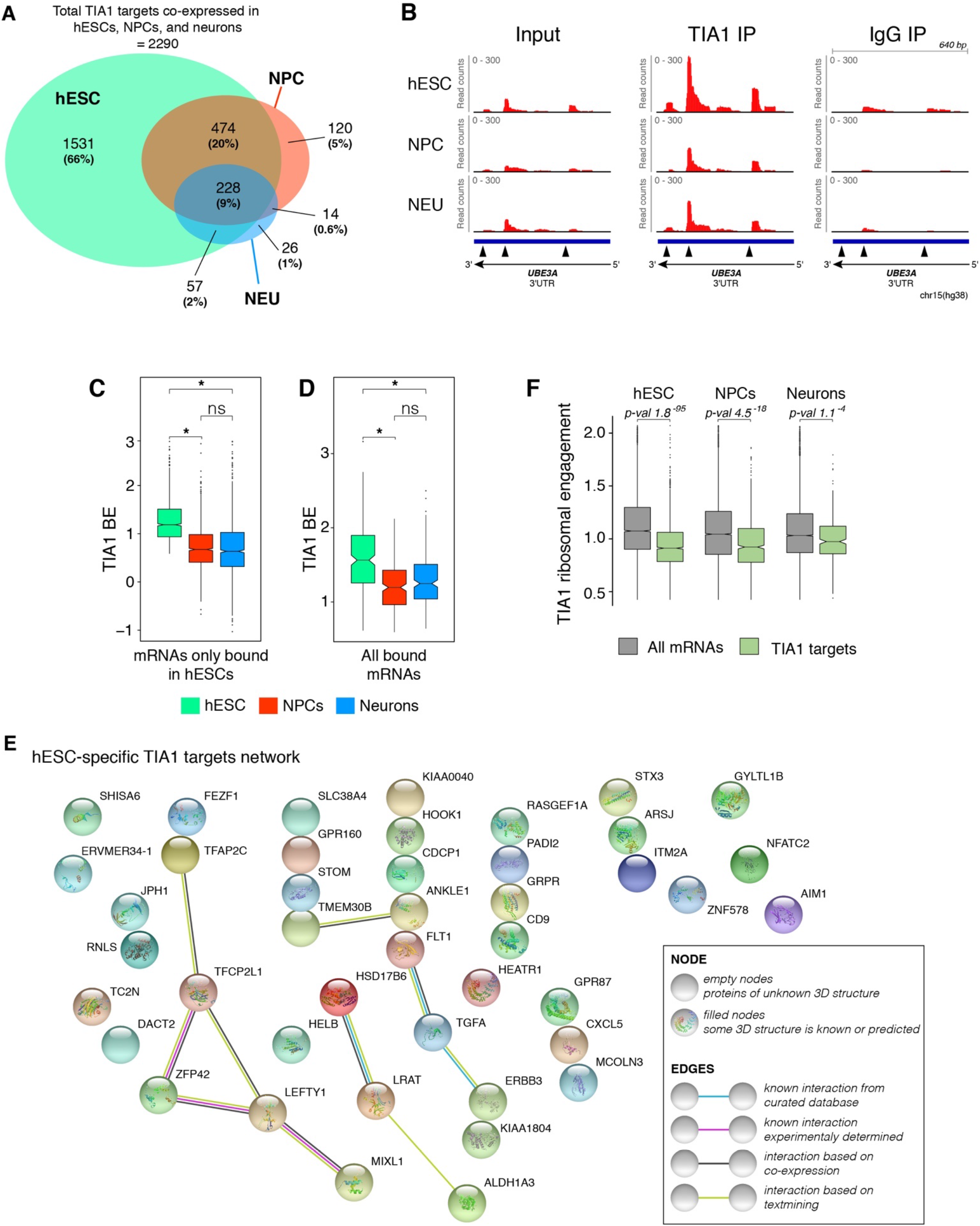
TIA1 network of target mRNAs decreases during human neurodevelopment and potential for translation regulation. A, Venn diagram showing overlap of TIA1 mRNA targets that are expressed in all cell types. B, density plots showing reads mapped to the polyadenylation sites in the 3ʹUTR of the TIA1 mRNA target UBE3A. Note the higher enrichment of reads in the TIA1 IP relative to the input and IgG samples, and the decrease in reads density in NPCs and neurons relative to hESCs. C, BE levels of TIA1 mRNA targets only bound in hESCs and expressed in all cell types (* p-val < 2.2^−16^. p values calculated using paired t-test). D, BE levels of TIA1 mRNA targets bound in all cell types. E, protein-protein interaction network map of hESC-specific TIA1 targets showing low functional association between genes. F, ribosomal engagement levels of TIA1 mRNA targets (green) relative to all detected genes (grey) in the three cell types. For C, D, and F the p-values were calculated using paired t-tests.

Finally, we investigated the potential cell-type specific functions of TIA1 mRNA binding. We looked at the genes expressed in all cell types that were at the bottom 20 percentile in the other two cell types. Using this threshold, we found that out of the 2,400 hESC TIA1 targets, only 47 are hESC specific (*i.e.* not expressed in NPCs or neurons) (Figure 5E). No molecular nor biological functions are enriched for these 47 genes, and protein-protein interaction network analyses only show a loose relationship between these genes (Figure 5E). Interestingly, we found no TIA1 targets exclusively expressed in NPCs and only *TMEM196* as a neuron-specific target gene (Supplementary table 4). Altogether, these results indicate that TIA1 plays a major role in the first stages of human neurodevelopment where most of its targeted mRNAs in hESCs continue to be expressed in NPCs and neurons but are no longer targeted by TIA1.

### TIA1 may regulate global translation during neurodevelopment and ASD

Since TIA1 has been shown to be a translational regulator in hESC [17], we tested whether we could detect changes in translational activity of TIA1 bound mRNAs during neurodevelopment. We re-analyzed an existing dataset of parallel TRAP-seq (*translating ribosome affinity purification*-seq) and RNA-seq that quantified changes in ribosomal engagement, hence translation regulation, transcriptome-wide in similar conditions of *in vitro* neurodevelopment [28]. Interestingly, we found that TIA1 mRNA targets in all cell types had, on average, significantly lower ribosomal engagement than transcripts not bound by TIA1, indicating that many of these transcripts could be translationally repressed by TIA1 (Figure 5F) at the ribosomal engagement level. This result is consistent with the role of TIA1 in repressing translation by recruiting mRNAs to translationally inactive ribonucleotide granules [11].

## Discussion

Here we investigated the roles of the RBP TIA1 during typical neurodevelopment by determining its network of targeted mRNAs in a hESC-derived neuronal developmental model. We performed TIA1 RIP-seq on multiple replicates of hESC, - derived NPCs and cortical neurons and calculated its BE to RNAs transcriptome-wide. We also deployed an isogenic IgG antibody as a negative control and to set the minimal BE threshold for target-mRNA calling. With this experimental set-up, we identified TIA1 target mRNAs with high confidence while minimizing false-positive calls, yielding 2,400 TIA1 targets in hESCs, 845 in NPCs, and 330 in neurons. Interestingly, these numbers of target mRNAs represent 18% of hESCs transcriptome, but only 7% and 3% of the transcriptome in NPCs and neurons, respectively, even though both TIA1 and the vast majority of the target mRNAs were expressed in all three cell types. We found only 47 TIA1 targets in hESCs that are not expressed in either NPCs or neurons. Additionally, the remaining TIA1 target mRNAs in NPCs and neurons had significantly decreased BE relative to hESCs. The decrease in the number of TIA1 target mRNAs and reduction of BE upon differentiation indicate that TIA1 dynamically regulates mRNA metabolism in the early stages of human neurodevelopment.

The underlying causes of the observed decrease in TIA1 binding during neurodevelopment are still unclear. One possibility is that the changes in the TIA1 mRNA network may be due to post-translational modifications to the protein, which would not be without precedent. For example, CPEB3 is an RBP that regulates mRNA translation in neurons and, like TIA1, CPEB3 is a prion-like protein that can self-aggregate via a prion-like domain [29]. Post-translational modifications such as sumoylation and ubiquitination have been shown to alter both the activity and aggregation of CPEB3 in neurons [30, 31]. Phosphorylation of TIA1 has also been reported during apoptosis [32]. However, changes in post-translational modifications of TIA1 between cell types have not been investigated.

The decrease in TIA1 BE in neurons could also be due to structural changes in the mRNA targets themselves, or the expression of other RBPs or miRNAs impacting TIA1 binding. Post-transcriptional regulation typically involves multiple regulators and is determined by the local concentrations of these various regulators in the cellular environment. Changes in the 3ʹUTR of TIA1 transcripts during differentiation could alter the balance between regulators and affect TIA1 binding. CLIP-seq experiments in both B-cells and HEK293s have shown that many TIA1 targets have multiple TIA1 binding sites within their introns or 3ʹUTRs and that TIA1 BE is directly proportional to the number of binding sites present [20, 27]. A loss of TIA1 binding sites, or the inaccessibility of existing sites, could explain the decrease in TIA1 BE. Additionally, some studies have shown that binding sites closer to the 3ʹ-end of the 3ʹUTR have higher BE than sites within the center of UTRs [33]. Lengthening of the 3′UTR by the alternative use of more distal polyadenylation sites, a feature of neurodevelopment [28, 34], could therefore position the TIA1 binding sites towards the center of the 3′UTRs in NPCs and neurons, making them less accessible.

Another possible explanation for the decrease in TIA1 binding in neurons is an increase in the abundance of other regulatory molecules that share a binding site with TIA1. At higher abundances, these factors can outcompete TIA1 for binding to transcripts changing their regulatory profile as has been previously shown [35, 36]. The Hu family of RBPs have repeatedly been implicated as competitors of TIA1, and three of the four members of this family (HuB, HuC, and HuD; gene names *ELAVL2*, *ELAVL3*, and *ELAVL4*, respectively) are expressed exclusively in a neuronal context [17, 36, 37]. Our attempts to test this hypothesis by performing RIP of Hu proteins have not been successful because the antibodies are not sufficiently specific to bind only one individual Hu protein. RIP-seq experiments to address this question will require the insertion of epitope tags to the Hu proteins, ideally in the endogenous locus using a knock-in approach to cause minimal effect on protein stoichiometry in the cell. Another RBP of interest that may compete with TIA1 is FMRP. FMRP is highly expressed in neurons and has previously been shown to regulate several transcripts that we identified in our TIA1 RIP-seq data set, including *UBE3A* [38].

Together, our results provide important insights into the network of TIA1 target mRNAs during neurodevelopment and its roles in neuronal differentiation. TIA1 appears to regulate neurodevelopmentally relevant transcripts in the early stages of development, and the network of regulated mRNAs decreases as cells differentiate to form neurons, despite the continued presence of TIA1 in the cells. This decrease in TIA1 binding may play an important role in regulating the transition from stem cells and progenitor cells to functional mature neurons.

## Acknowledgments

We thank The Centre for Applied Genomics for RNA-seq.

## Declarations

### Ethics approval

Pluripotent stem cell work was approved by the Canadian Institutes of Health Research - Stem Cell Oversight Committee and The Hospital for Sick Children Research Ethics Board.

### Authors’ contributions

Conceptualization, L.P.B., D.C.R., and J.E.; Investigation, L.P.B., M.M., K.E.Y., W.W., and A.P.; Software, M.M.; Writing – Original Draft, L.P.B., D.C.R., and J.E.; Writing – Review & Editing, L.P.B., D.C.R. M.M., K.E.Y., W.W., A.P., M.D.W., and J.E.; Visualization, L.P.B., M.M., and D.C.R.; Supervision, M.D.W., D.C.R., and J.E.; Funding Acquisition, M.D.W., and J.E.

---

**Supplementary figure 1.**
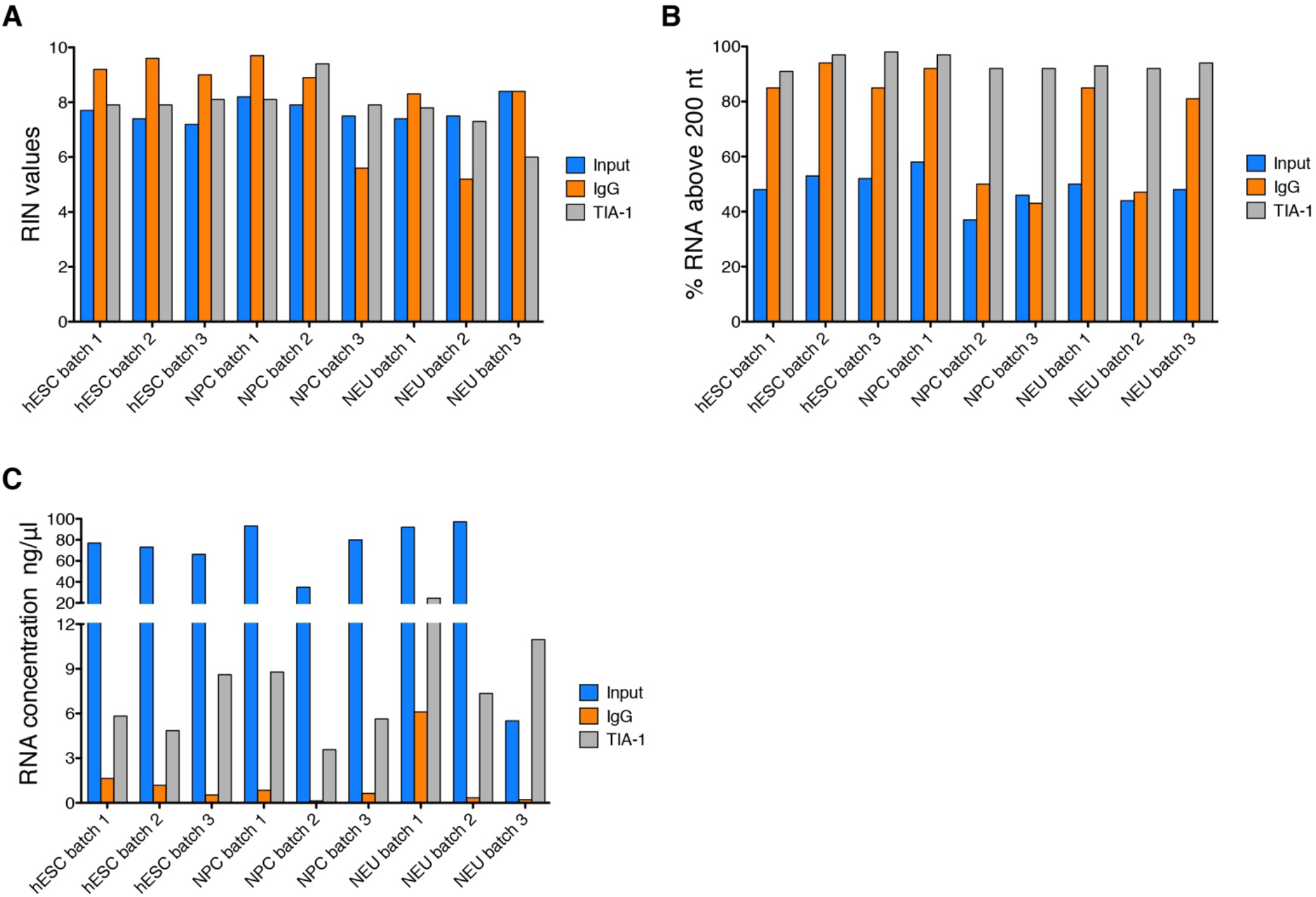
Quality control of RNA samples used for sequencing. A, RNA Integrity Numbers (RIN) from all RNA samples show that all input samples have RIN values above 7. B, percentage of RNA molecules larger than 200 nucleotides for each sample. C, RNA concentration of all samples show that IgG samples had low yields in RNA recovery.

## Methods

### Cell Culture and cortical neuronal differentiation

H9 hESCs obtained from the National Stem Cell Bank (WiCell) were cultured with mTeSR1 culture medium (STEMCELL Technologies) on Becton Dickinson hESC-qualified matrigel. Pluripotent stem cell work was approved by the Canadian Institutes of Health Research Stem Cell Oversight Committee and The Hospital for Sick Children Research Ethics Board.

An embryoid body (EB)-based method was used for neuronal induction of NPCs as described previously [39]. hESCs were cultured as cellular aggregates in low-attachment dishes in N2 media containing laminin (1 μl/ml) with 10 μM SB431542, 2 μM DSM and 1x penicillin-streptomycin changed daily. After 7 days, EBs were plated on poly-L-ornithine + laminin coated dishes and grown in N2 media + laminin (1 μl/ml). After 7 days, neural rosettes were manually picked and transferred to poly-L-ornithine+laminin coated wells. After 7 days, neural rosettes were picked a second time, digested with Accutase and plated on poly-L-ornithine + laminin coated wells. Resulting neural precursor cells were grown as a monolayer and split every 5-7 days in NPC media (DMEM/F12, N2, B27, 1x non-essential amino acid (NEAA), 2 μg/ml Heparin, 1 μg/ml laminin). To generate neurons, NPCs were plated on poly-L-ornithine + laminin-coated plates at a density of 10^6^ cells per 10 cm dish and cultured for 4 weeks in neural differentiation medium (Neurobasal, N2, B27, 1 μg/ml laminin, 1x penicillin-streptomycin, 10ng/ml IGF, 10ng/ml BDNF, 10ng/ml GDNF, 200 μM ascorbic acid, and 10 μM cAMP).

### siRNA knockdown of TIA1 in hESCs

hESCs were transfected with Silencer select siRNAs (ThermoFisher) to knockdown TIA1 (# s14133 and s14132) following manufacturer’s instructions. Cells were subjected to two consecutive transfections of siRNAs at two-day intervals in the week leading up to protein harvest.

### Western blot

Cells were washed in ice-cold PBS and total protein extracted in radioimmune precipitation assay (RIPA) buffer (25 mM Tris-HCl, pH 7.6, 150 mM NaCl, 1% Nonidet P-40, 1% sodium deoxycholate, and 0.1% SDS). Equivalent protein mass was loaded on SDS-PAGE and transferred to Hybond ECL (GE HealthCare) nitrocellulose membrane. Membranes were incubated with TIA1 and Actin B antibodies (Abcam #ab40693 and Sigma #A5441, respectively) in 5% milk in PBS+Tween 20 (0.05%). Near-Infra Red-conjugated secondary antibodies (LI-COR) were used and membranes scanned using LI-COR Odyssey CLx scanner according to manufacturer’s instructions. Acquired images were analyzed using ImageStudio v5.2.5.

### RNA Immunoprecipitation – RIP

For each RIP lysate hESCs or NPCs were seeded at a density of 1×10^7^ cells per plate, on six 10 cm plates, and grown overnight prior to lysate collection. To collect neuronal samples, NPCs were seeded on 10 cm plates and maintained in neuronal differentiation media for 4-weeks prior to lysate collection. Lysate collection and RNA immunoprecipitations were performed using the Magna RIP™ RNA-Binding Protein Immunoprecipitation Kit (Millipore #17-700) following manufacturer’s instructions. Briefly, cells were washed twice in ice-cold PBS and lysed in RIP Lysis Buffer complemented with Protease and RNase Inhibitor Cocktails. Prior to immunoprecipitation, magnetic protein A/G beads were prepared with 5 μg of rabbit anti-TIA1 (Abcam - #ab40693) or normal rabbit IgG antibodies (from Magna RIP kit). The RIP lysates were centrifuged at 12.000 g for 10 minutes at 4°C. The lysate was then split into three fractions: 10 μL was set aside as an input sample and frozen at −80°C until the RNA purification step, 100 μL of RIP lysate was added to the tube containing the TIA1 bound beads, and the IgG bound beads. The RIP lysates were incubated with the antibody-bead complexes overnight at 4°C. Total RNA was isolated and purified from the input lysates and the immunoprecipitation samples using phenol/chloroform and resuspended in 10 μL of RNase-free water. Quality control of the RIP samples was assessed by qRT-PCR (see below). RNA samples were sent to The Centre for Applied Genomics (TCAG) for bioanalyzer analysis to assess the quality of the RNA.

### qRT-PCR (RIP-qRT-PCR)

RNA was reversed transcribed to cDNA using SuperScript III reverse transcriptase, and random hexamer primers. qRT-PCR was performed using SYBR Select PCR master mix on a ViiA7 Real-Time PCR System (both from ThermoFisher). Fold changes were calculated by the 2^−(ΔΔCt)^ method, and 18s rRNA as a normalizing control. Primers used were: *MECP2* forward: GCUCUGCUGGGAAGUAUGAUG, reverse: TTTGGGCTTCTTAGGTGGTTT; *UBE3A* forward: GTTCTGATTAGGGAGTTCTGGG, reverse: TCCTTTGGCATACGTGATGG; and *18S* forward: GATGGGCGGCGGAAAATAG, reverse: GCGTGGATTCTGCATAATGGT. Technical duplicates and biological replicates were combined to calculate an average fold change. For comparison of mRNA levels between input and IP samples, results are displayed as a percentage of input.

### QuantSeq 3’ mRNA library preparation

Sequencing libraries were prepared using the Agilent Bravo automation system (Agilent Technologies, Santa Clara) using a previously established protocol: (https://www.agilent.com/cs/library/applications/5991-8601EN_Auto_NGS_Application.pdf). Library preparation was done using the QuantSeq 3′ mRNA-Seq Library Prep Kit for Illumina (Lexogen #015.24), following manufacturer’s instructions. To minimize variability in the library preparation, 17 ng of RNA was used in the library preparations of all input and TIA1 IP samples. The RNA was spiked with SIRV Set 3 Iso Mix E0/ERCC controls (Lexogen 025.03) prior to the start of library prep. Due to low RNA levels in the IgG samples the maximum amount of 5 μL of RNA was used for each IgG sample. Each sample was dual indexed during the final library amplification using the Lexogen’s i5 6 nt Dual Indexing Add-on Kit (#047) to allow for differentiation of the samples after sequencing.

### Sequencing

The sequencing was then conducted by The Centre for Applied Genomics (TCAG) using the Illumina HiSeq 2500 and two lanes of the Rapid Run Mode flow cell. The samples were sequenced as single-ended reads, with a read length of 100 base pairs, at a sequencing depth of approximately 20 million reads for each sample.

### Quantification of polyadenylation (polyA) sites

The unique molecular identifiers were used to remove PCR duplicates. Reads were then aligned to the human genome hg38 using the STAR aligner [40], with default settings. Reads were assigned to previously mapped polyA sites [41]. This map is a recent polyA site database generated 3′READS from 20 human cell types, including hESCs, neural stem cells, and cerebellum. Each polyA site is a window of 30-nucleotides stretching from 20-nucleotides upstream of the polyA site to 10-nucleotides downstream of the polyA site to account for random priming, and wobbling of the location of the cleavage site [23]. Only reads that were overlapping with this 30-nucleotide window were assigned to polyA sites. Counts from all polyA sites within a gene were then combined to estimate gene abundance.

### Quality control of sequencing and quantifications

Raw gene counts were normalized to corresponding medians of each replicate and cell-type. Then, *log10* of normalized counts were plotted in a heatmap showing abundance of cell-type specific markers. Second, we used variance stabilizing transformation on raw gene counts (*vst* function in DESeq2 package). Then, PCA of transformed counts was used to validate clustering of samples from same assay and cell type (*plotPCA* in DESeq2 package).

### Reads visualization

Reads density distributions in the human genome were visualized using STAR aligned reads with IGV version 2.3.69 [42].

### Estimate of TIA1 binding efficiency

TIA1 binding efficiency was defined as a ratio of TIA1 IP and Input mRNA abundances, corresponding to a fraction of TIA1 bound mRNAs, as done previously [12][25]. To eliminate gene specific IP bias, enrichment of TIA1 versus IgG IP was used. First, for TIA1 IP samples, library size factors in DESeq2 were scaled by 1.5 relative to IgG samples (manually defined: sizeFactors(dds) = 1.5*estimateSizeFactorsForMatrix(counts)) for all cell-types. This scaling factor conservatively accounts for consistently higher RNA yields in TIA1 over IgG IP. In our hands, RNA yield was at least 4 times higher in TIA1 than IgG IP, performed from the same number of cells. Then, genes were selected for next step when TIA1 over IgG enrichment was higher than 1.5 (equivalent to ≥ 0.58 in *log2* scale) and adjusted p-values ≤ 0.05, estimated in DESeq2 with size factors defined above and the design: ~ replicate + assay. Assay corresponds to TIA1 IP, IgG IP and Input samples. Finally, TIA1 mRNA targets were defined from selected genes, when TIA1 IP over Input enrichment was higher than 1.5 (equivalent to ≥ 0.58 in *log2* scale) and adjusted p-values ≤ 0.05, estimated in DESeq2 with design: ~ replicate + assay. Replicate factor in the DESeq2 design is used to reduce batch noise by matching TIA1 IP, IgG IP and Input samples from the same lysate. Roughly, this approach estimates an average binding efficiency (BE) and its accuracy for each transcript across all three biological replicates.

### Ribosomal engagement (RE) of TIA1 mRNA targets

RE of all genes was downloaded from GEO: GSE123753 [28]. Then, RE was compared between two groups: all mRNAs versus cell-type specific targets.

### Gene ontology and interaction of TIA1 targets

Gene ontology (GO) analysis was done using g:profiler and default parameters [43]. Genes that were identified as TIA1 targets in hESCs were analyzed for enrichment in terms associated with biological processes, against the background of all genes detected in hESCs. Enriched terms were defined using a Benjamini-Hochberg false discovery rate < 0.05. The resulting GO terms were then used to construct an enrichment map in Cytoscape [44] to identify the key cellular functions regulated by TIA1 in hESCs. During construction of the enrichment maps, unclustered GO-terms were removed. Cluster names were edited manually for readability. Gene interaction network map of hESC-specific TIA1 targets was done using String version 11.0 using default parameters [45].

### Statistics

All datasets and statistical tests were done using R software. For comparison of TIA1 binding efficiency of different transcripts within the same samples statistical analysis was performed using two-tailed student’s t-test. For comparison of ribosomal engagement of the same transcripts in different cell types statistical analysis was done using a paired t-test. For RIP-seq data analysis, the P-values were adjusted using the Benjamini-Hochberg method. In all cases, fold-change differences with a p-value ≤ 0.05 were considered significant.

